# Exact Tests of Zero Variance Component in Presence of Multiple Variance Components with Application to Longitudinal Microbiome Study

**DOI:** 10.1101/281246

**Authors:** Jing Zhai, Kenneth Knox, Homer L. Twigg, Hua Zhou, Jin J. Zhou

## Abstract

In the metagenomics studies, testing the association of microbiome composition and clinical conditions translates to testing the nullity of variance components. Computationally efficient score tests have been the major tools. But they can only apply to the null hypothesis with a single variance component and when sample sizes are large. Therefore, they are not applicable to longitudinal microbiome studies. In this paper, we propose exact tests (score test, likelihood ratio test, and restricted likelihood ratio test) to solve the problems of (1) testing the association of the overall microbiome composition in a longitudinal design and (2) detecting the association of one specific microbiome cluster while adjusting for the effects from related clusters. Our approach combines the exact tests for null hypothesis with a single variance component with a strategy of reducing multiple variance components to a single one. Simulation studies demonstrate that our method has correct type I error rate and superior power compared to existing methods at small sample sizes and weak signals. Finally, we apply our method to a longitudinal pulmonary microbiome study of human immunodeficiency virus (HIV) infected patients and reveal two interesting genera *Prevotella* and *Veillonella* associated with forced vital capacity. Our findings shed lights on the impact of lung microbiome to HIV complexities. The method is implemented in the open source, high-performance computing language Julia and is freely available at https://github.com/JingZhai63/VCmicrobiome.

## 1. Introduction

Technology advances led to a much deeper understanding of microbes and their link to human health (Eckburg et al., 2005; Haas et al., 2011; Hodkinson and Grice, 2015; Kuleshov et al., 2016; Wang and Jia, 2016). In particular, for the pulmonary microbiome, Rogers et al. (2010) hypothesized that microbial lung community might exist and can be considered as a unique, distinct pathogenic entity. The culture-independent microbial detection method, 16S ribosomal RNA (rRNA) gene sequencing, demonstrated the existence of pulmonary microbiome, both in healthy (Erb-Downward et al., 2011; Morris et al., 2013; Twigg et al., 2013) and disease populations (Lozupone et al., 2013; Zemanick et al., 2011).

The Lung HIV Microbiome Project (LHMP) (Grubb et al., 2006) studies the respiratory microbiome of HIV-infected patients and how the highly active antiretroviral therapy (HAART) may alter its construction. A longitudinal cohort of HIV-infected subjects were collected before and up to three years after starting HAART. For a quantitative phenotype in a longitudinal design, one can propose

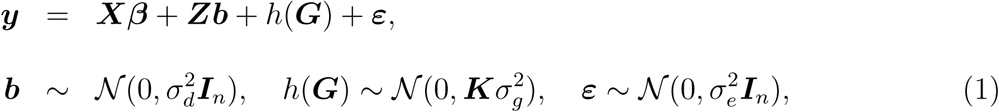

where ***y*, *X*, *G*** and ***ε*** are the vertically stacked vectors/matrices of ***y***_*i*_, ***X***_*i*_, ***G***_*i*_ and ***ε***_*i*_. ***y***_*i*_ is a vector of *n*_*i*_ repeated measures of a quantitative phenotype for individual *i*. ***X***_*i*_ is the *n*_*i*_ *× p* covariate matrix. ***G***_*i*_ is an *n*_*i*_ *× u* Operational Taxonomic Unit (OTU) abundance matrix for individual *i* (*u* is the total number of OTUs). These OTUs are related by a known phylogenetic tree. ***ε***_*i*_ is an *n*_*i*_ *×* 1 vector of the random error. ***Z***_*i*_ = **1**_*n_i_*_ links the random intercept *b*_*i*_ to ***y***_*i*_. ***Z*** is a block diagonal matrix with ***Z***_*i*_ on its diagonal. ***β*** is a *p ×* 1 vector of fixed effects. ***b*** = (*b*_*i*_) is the subject-specific random effects. ***K*** is a kernel matrix capturing distances between individuals, e.g., the UniFrac distance (Lozupone and Knight, 2005) or the Bray-Curtis dissimilarity (Bray and Curtis, 1957) (Web Appendix A). Therefore,

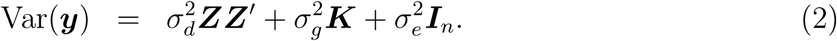

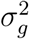 is the phenotypic variance explained by microbiome. 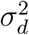 is the phenotypic variance due to the correlation of repeated measurements, and 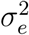 is the within-subject variance that cannot be explained by microbiome and repeated measurements. To detect the overall microbiome association is to test 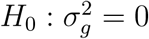 versus 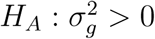. When 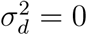 model (1) reduces to the microbiome regression-based kernel association test (`MiRKAT`) (Zhao et al., 2015; Chen et al., 2016; Zhan et al., 2017). In this example, the extra variance component 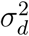 is necessary to capture the correlation between repeated measurements.

After the overall association is identified, localization of the signal to a specific component of the microbial community is essential for downstream mechanistic studies and drug discoveries. For instance, Jangi et al. (2016) found that multiple sclerosis patients had significantly increased abundance of the phylum *Euryarchaeota*. However, such fine cluster effects can be tagged by other correlated microbial in the community (Gilbert et al., 2016), leading to false positive discoveries. To detect association from specific taxonomic clusters, distances and kernel matrices can be formulated using abundances and tree information only from a specific cluster. Overall microbiome effects are then partitioned into different clusters at the same taxonomic level. That is

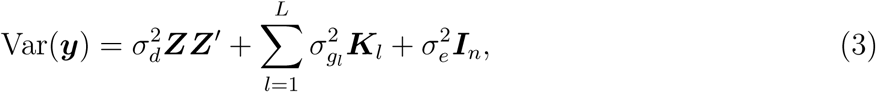

where *L* is the total number of clusters. We are now interested in testing specific taxonomic cluster effects: 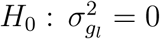 versus 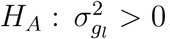.

Current methods for testing null variance component in model (2) and model (3) are based on either asymptotics or parametric bootstrap. Under the assumption that the response variable vector can be partitioned into independent and identically distributed (IID) subvectors and the number of independent subvectors is sufficient, asymptotic null distribution of the likelihood ratio, Wald, and score tests are available (Self and Liang, 1987; Stram and Lee, 1994; Silvapulle and Sen, 2011). However, the asymptotic approximation deteriorates when the data are highly correlated without a sufficient number of independent blocks. When *m* = 1, Crainiceanu and Ruppert (2004) developed a computational procedure for obtaining the approximate finite-sample distribution of the likelihood ratio and restricted likelihood ratio test statistics. Greven et al. (2008) provided a pseudolikelihood-heuristic extension of this method to the *m* > 1 situation. Later Drikvandi et al. (2013) proposed a permutation test that does not depend on the distribution of the random effects and errors except for their mean and variance and can be applied to the *m* > 1 situation. However, the permutation test is computationally prohibitive for high dimensional tests. Qu et al. (2013) proposed a test statistic that is the weighted sum of the scores from the profile likelihood. Their method considered testing a subset of the variance components to be zero. When *m* = 1, Qu et al. (2013)’s method is exact; when *m* > 1, their test relies on asymptotic theory. Besides, score-based tests can be less powerful than the likelihood ratio tests, especially when sample sizes are limited as in most of the sequencing studies. Saville and Herring (2009) developed yet another type of test based on the Bayes factors using Laplace approximation. It cannot be easily extended to multiple random effects, and relies on the subjective choice of the prior distribution of parameters. Others have suggested procedures based on Markov chain Monte Carlo methods (Chen and Dunson, 2003; Kinney and Dunson, 2007), but they can be time-consuming, especially when the number of random effects is large.

In this article, we propose methods of performing exact likelihood ratio test (`eLRT`), exact restricted likelihood ratio test (`eRLRT`), and exact score test (`eScore`) of a variance component being zero for the finite sample. Our approach combines the corresponding exact tests for the *m* = 1 case with a strategy of reducing the *m* > 1 case to the *m* = 1 case (Ofversten, 1993; Christensen, 1996). Our method is the first one that provides `eLRT`, `eRLRT`, and `eScore` for testing zero variance component when multiple variance components are present (*m* > 1).

## 2. Methods

### 2.1 *Exact tests with one variance component under H*_0_

We briefly review the three exact tests, `eLRT`, `eRLRT`, and `eScore`, for testing 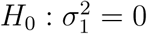 in model

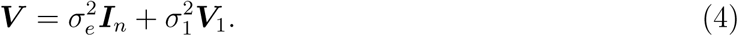

Note the change of notation for general modeling. In the previous motivating microbiome example, 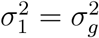 and ***V***_1_ = ***K***, the kernel matrix calculated from microbiome abundances. A slight extension allows for testing the more general case 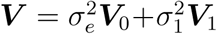 where ***V***_0_ *∈* ℝ^*n×n*^ is a known positive semidefinite matrix. Let *t* = rank(***V***_0_). Given the thin eigen-decomposition ***V***_0_ = ***UDU*** ^*′*^, define ***T*** = ***D***^*–*1*/*2^***U*** ^*′*^ *∈* ℝ^*t×n*^. Then 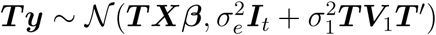 and the `eLRT` and `eRLRT` (Crainiceanu and Ruppert, 2004) or the `eScore` test (Zhou et al., 2016) can be applied to ***T y***.

Let 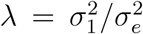 be the signal-to-noise ratio, *s* = rank(***X***), and write the covariance as 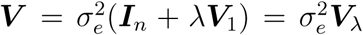. The model parameters are 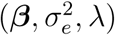. Testing 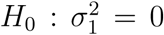 is equivalent to testing *H*_0_: *λ* = 0. The log-likelihood function is 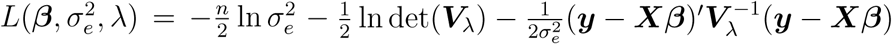. The likelihood ratio test (LRT) statistic is

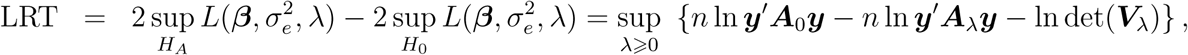

where ***P*_*X*_** = ***X***(***X***^*′*^***X***)^*–*1^***X***^*′*^ is the projection matrix onto the column space *𝒸*(***X***), ***A***_0_ = ***I*** *–* ***P*_*X*_** and **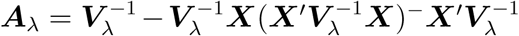**. Let *{ξ*_1_, *…, ξ*_*l*_ *}* be the positive eigenvalues of ***V***_1_ and *{µ*_1_, *…, µ*_*k*_*}* the positive eigenvalues of ***A***_0_***V***_1_***A***_0_. Then

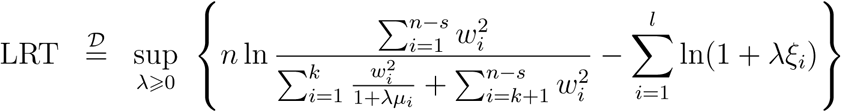

where, under the null, *w*_*i*_ are (*n – s*) independent standard normals. Under the alternative, *w*_*i*_ *∼ 𝒩* (0, 1 + *λµ*_*i*_) for *i* = 1, *…, k*, *w*_*i*_ *∼ 𝒩* (0, 1) for *i* = *k* + 1, *…, n – s*, and they are jointly independent. The null distribution can be obtained from computer simulation (Web Appendix B, Algorithm 1). A computationally efficient *χ*^2^ approximation algorithm is given in the Supplementary Material (Web Appendix B, Algorithm 4). The same derivation can be carried out for the `eRLRT`, in which case

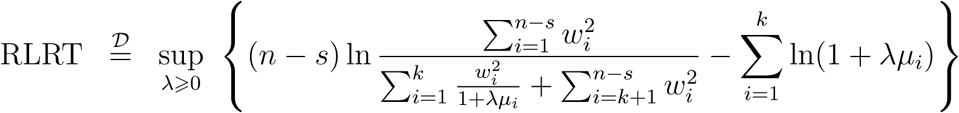

The null distribution for `eRLRT` can be obtained from computer simulation (Web Appendix B, Algorithm 2). Algorithms 1 and 2 in Web Appendix B contain a univariate optimization for each simulated point from the null distribution and can be computationally intensive for obtaining extremely small *p*-values. To further reduce computational burden, we adopt the Satterthwaite method to approximate the null distributions (Zhou et al., 2016).

For `eScore`, it is easier to work with the original parameterization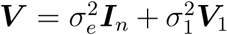. The (Rao) score statistic is based on 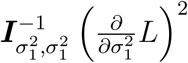, where the information matrix and score are evaluated at the maximum likelihood estimator (MLE) under the null. The resultant test rejects the null when

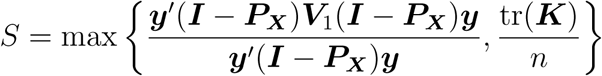

is large. Let *{µ*_1_, *…, µ*_*k*_*}* be the positive eigenvalues of (***I*** *–* ***P*_*X*_**)***V***_1_(***I*** *–* ***P*_*X*_**). Then

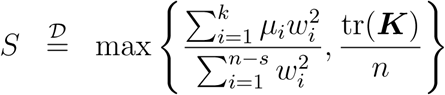

where *w*_*i*_ are *n – s* independent standard normals. The null distribution can be obtained from computer simulation as shown in Algorithm 3 in Web Appendix B or inverting the characteristic function (Zhou et al., 2016). Both options, simulation (Web Appendix B, Algorithms 1, 2, and Algorithm 3) and approximation of null distribution, are available in our program, https://github.com/JingZhai63/VCmicrobiome.

### 2.2 *Exact tests with more than one variance components under H*_0_

In this section we consider the situation when ***Y*** *∼ 𝒩* (***Xβ***, ***V***) with 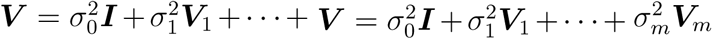 and *m* < 1. We are interested in testing 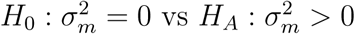. Our strategy is to reduce the problem to the *m* = 1 case in section 2.

We first obtain an orthonormal basis (***Q***_0_, ***Q***_1_, *…*, ***Q***_*m*_, ***Q***_*m*+1_) of ℝ^*n*^ such that ***Q***_0_ is an orthonormal basis of *𝒞*(***X***), ***Q***_1_ is an orthonormal basis of *𝒞*(***X***, ***V***_1_) – *𝒞*(***X***), ***Q***_*i*_ is an orthonormal basis of *𝒞*(***X***, ***V***_1_, *…*, ***V***_*i*_) – *𝒞*(***X***, ***V***_1_, *…*, ***V***_*i-*1_) for *i* = 2, *…, m*, and ***Q***_*m*+1_ is an orthonormal basis of ℝ*^n^ – 𝒞*(***X***, ***V***_1_, *…*, ***V***_*m*_). Denote their corresponding ranks by *r*_0_, *…, r*_*m*+1_. If *r*_*m*_ *>* 0, that is *𝒞* (***X***, ***V***_1_, *…*, ***V***_*m–*1_) < ⊊ *𝒞* (***X***, ***V***_1_, *…*, ***V***_*m–*1_, ***V***_*m*_), then 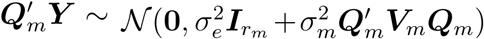 and `eLRT`, `eRLRT` and `eScore` can be applied to 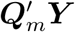. The order of ***V***_1_, *…*, ***V***_*m*_ does not matter. If *r*_*m*_ = 0, that is *𝒞* (***X***, ***V***_1_, *…*, ***V***_*m–*1_) = *𝒞* (***X***, ***V***_1_, *…*, ***V***_*m*_), we construct a test based on the transformed data 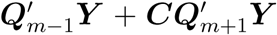. Without loss of generality we assume ***Q***_*m–*1_ is nontrivial. If *r*_*m–*1_ = 0, we use ***Q***_*m–*2_ and so on. We consider the following cases:

1. If 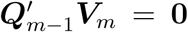 when *𝒞* (***V***_*m*_) *⊂ 𝒞* (***X***, ***V***_1_, *…*, ***V***_*m–*2_), then this test cannot be performed. Shifting the order of ***X***, ***V***_1_, *…*, ***V***_*m–*1_ might solve the issue.
2. If 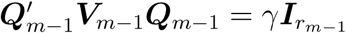 and *γ* ≠ 0, then

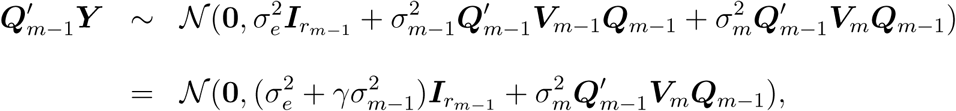

which is the case (4). `eLRT`, `eRLRT` and `eScore` can be applied without using the 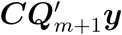 piece.
3. If 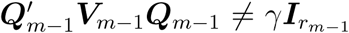 then the test requires the 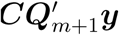 term. 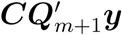 has distribution 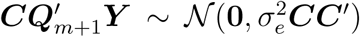. Since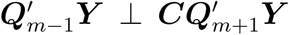 we pick ***C*** such that

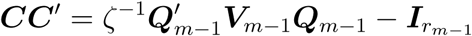

where the scalar *ζ* is chosen such that 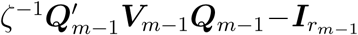 is positive semidefinite. Let 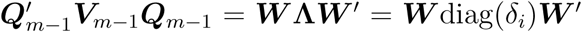 be the eigen-decomposition, *ζ* be the smallest positive eigenvalue, and ***C*** = ***W*** diag 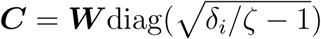. Then the transformed data

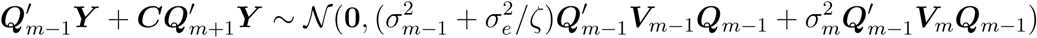

and the test for case (2.1) can be applied. A larger *ζ* leads to a higher signal-to-noise ratio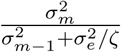 and thus a more powerful test. Finally we test 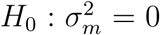 using `eLRT`, `eRLRT` or `eScore` test on the transformed data,

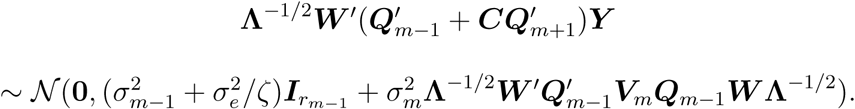

We note that if in some applications that matrices have high or full rank, consuming most or all available degrees of freedom after above reduction strategy. One could proceed with a low rank approximation. For example, if *m* = 2 and ***V***_1_ has high or full rank, one could find rank *r***_*v1*_** approximation of ***V***_1_ as follows: let *r***_*K*_** = rank(***V***_2_), ***Q***_0_ is an orthonormal basis of *𝒞* (***X***), and *r*_0_ = rank(***Q***_0_). A rank 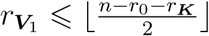 approximation of ***V***_1_ is suffice to perform testing. Details can be found in the software’s documentation (http://vcmicrobiomejl.readthedocs.io/en/latest/).

## 3. Simulation

We evaluate the performance of the exact tests for longitudinal microbiome study in three simulation scenarios (Table 1).

**Table 1.** *Simulation configurations. For all simulations*,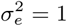 *and* 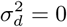 *when number of repeats (# Repeat)* 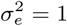 *and* 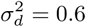 *when number of repeats >* 1. *There are* 2964 *OTUs presented in simulated count data. A phylogenetic tree generated using the real pulmonary microbiome data is used for kernel calculation and phenotype simulations.* ***K***_*W*_: *weighted UniFrac kernel;* ***K***_*U*_ *: unweighted UniFrac kernel;* ***K***_*V AW*_ *: variance adjusted weighted UniFrac kernel; **K**α: generalized UniFrac kernels with α* = 0 *and* 0.5

| Sample size | Kernel type | Clustering | # Repeat | $\sigma_g^2$ | Method |
| --- | --- | --- | --- | --- | --- |
| 100 | $\mathbf{K}_W, \mathbf{K}_U, \mathbf{K}_{VAW}, \mathbf{K}_\alpha$ | None | 2 | 0 - 1.5 | eRLRT eScore |
| 100 | $\mathbf{K}_W, \mathbf{K}_U, \mathbf{K}_{VAW}, \mathbf{K}_\alpha$ | None | 1 | 0 - 1.5 | eRLRT eLRT eScore |

**Scenario 2: Localizing fine microbiome cluster effects**
| Sample size | Kernel type | Clustering | # Repeat | $\sigma_g^2$ | Method |
| --- | --- | --- | --- | --- | --- |
| 100 | $\mathbf{K}_W$ | Yes | 2 | 0 - 1.5 | eRLRT eScore |
| 100 | $\mathbf{K}_W$ | Yes | 1 | 0 - 1.5 | eRLRT eScore |

**Scenario 3: Comparing to existing methods**
| Sample size | Kernel type | Clustering | # Repeat | $\sigma_g^2$ | Method |
| --- | --- | --- | --- | --- | --- |
| 20, 30, 50, 100 | $\mathbf{K}_W$ | None | 2 | 0 - 1.5 | eRLRT eScore<br>LinScore |
| 20, 30, 50, 100 | $\mathbf{K}_W$ | None | 1 | 0 - 1.5 | eRLRT eLRT<br>eScore LinScore<br>MiRKAT |

Longitudinal microbiome count data with 2 repeated measurements are simulated using the R package ZIBR (Zero-Inflated Beta Random Effect model) (Chen and Li, 2016). To mimic features of real microbiome datasets, the phylogentic structure and average count information are extracted from the real HIV longitudinal pulmonary microbiome data. This microbiome dataset contains 30 samples, each with 2 to 4 repeated measurements: baseline, 4 weeks, 1 year and 3 years (Twigg III et al., 2016). OTU alignment at species level was produced by software Mothur (2009) (Schloss et al., 2009) and Basic Local Alignment Search Tool (BLAST, 1990; Altschul et al., 1990) in the Ribosomal Database Project (RDP) 16S database release 11.4 (Maidak et al., 1996). The phylogenetic tree at the OTU level is generated using the RDP classifier (Twigg III et al., 2016). We construct the higher taxon level, e.g., phylum, using the phylogenetic tree generator phyloT (2006) (Letunic and Bork, 2007, 2011) and NCBI database Taxonomy (1991) (Federhen, 2012). There are 2964 operational taxonomic units (OTUs) in total, 292 genera, and 24 phyla. Different distance measures are calculated using our Julia package PhylogeneticDistance.jl (2017). The definition of different distance measures and the details of simulation of microbiome abundances are provided in Web Appendix A and C.

Phenotypes are generated under three different scenarios. For all three scenarios, two covariates are included in the model. One of them is correlated with microbiome abundances. For individual *i*, *X*_1*i*_ *∼ 𝒩* (0, 1) and *X*_2*i*_ = *h*(***G***_*i*_)_*baseline*_ + *N* (0, 1). Their effects are *β*_1_ = *β*_2_ = 0.1. We set within-individual variance as 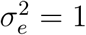. For longitudinal data simulation, between individual variance 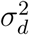 is set to 0.6. This corresponds to 60% of overall baseline phenotypic variance (Twigg III et al., 2016).

**Scenario 1: Testing overall microbiome effect.** Longitudinal responses are generated using model, 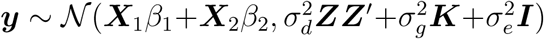, where 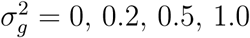 and 1.5. Wecompare the performance of five different distance measures: unweighted UniFrac (Lozupone and Knight, 2005), weighted UniFrac distance (Lozupone et al., 2007), variance adjusted weighted (VAW) UniFrac distance (Chang et al., 2011), and generalized UniFrac distance with parameter *α* = 0.0 and 0.5 (Chen et al., 2012).

**Scenario 2: Localizing fine microbiome cluster effects.** We cluster OTUs into 6 phyla, *Actinobacteria*, *Bacteroidetes*, *Fusobacteria*, *Proteobacteria*, *Firmicutes*, and *other*. We assume that only cluster *other*, *h*(***G***_1*i*_), has effects. That is 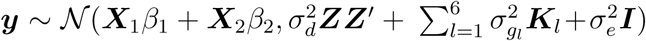 where 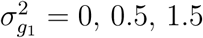 and 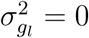 for *l* = 2, *…*, 6. Due to the correlation between phyla, marginal tests of 5 individual phylum may show false signal if we do not adjust for the effects of *h*(***G***_1*i*_). We present testing of variance components in a joint model has correct type I error.

**Scenario 3: Comparing with existing methods.** We compare our method with `MiRKAT` (Zhao et al., 2015) and `LinScore` (Qu et al., 2013). As `MiRKAT` can only be used for testing overall microbiome effects for cross-sectional designs, we first compare three methods when 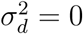. Responses are generated according to simulation scenario 1, where 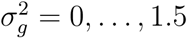.

In scenario 1 and 2, the sample size is fixed at *n* = 100. In scenario 3, we compare the performance of three methods under different sample sizes 20, 30, 50 and 100. The performance of five different kernels is compared in scenario 1. For scenarios 2 and 3, we focus on the weighted UniFrac distance kernel only, which has superior power in simulation scenario 1 than other kernels. 1000 Monte Carlo replicates are generated for all simulations and we use the nominal significance level 0.05 to evaluate type I error and power.

## 4. Results

### Simulation Results

**Scenario 1: Testing overall microbiome effect.** The type I error rate of `eRLRT`, `eLRT` and `eScore` tests with various distance kernel matrices using real longitudinal OTU count data are shown in Table 2. Figure 1 shows the power comparison with different kernels. In Figures 1*a* and 1*c*, five different kernels are constructed using OTU count data directly. In Figures 1*b* and 1*d*, OTU counts are summarized at the phylum level for kernel calculations. Figure 1 shows that kernel type greatly impacts the power. The weighted UniFrac kernel yields the highest power and the unweighted UniFrac kernel is the least powerful (Figures 1a and 1c). The pattern of the power increasing with effect size differs according to which taxon level count data are used to calculate the kernels. The power of five kernels became more similar among each other in Figures 1*b* and 1*d*. Moreover, the power of unweighted UniFrac kernel *K*_*UW*_, the least powerful one in Figures 1*a* and 1*c*, greatly improves in Figures 1*b* and 1*d*. This is because when the reads are summarized at the higher phylum level, the difference of abundance between each phylum is less notable. The less variation of abundance for each lineage, the more similar power for detecting microbiome association among each kernel types. As expected, reducing variance components leads to reduced degrees of freedom for association testing and the test is slightly less powerful in the longitudinal study compared to the cross-sectional study given the same effect strength.

**Table 2.** *Scenario 1: Type I error of `eLRT`, `eRLRT` and `eScore` for detecting overall microbiome effects. Five distance measures, weighted UniFrac kernel (****K***_*W*_ *), unweighted UniFrac kernel (****K***_*U*_ *), variance adjusted weighted UniFrac kernel (****K***_*V* *AW*_ *), and generalized UniFrac kernels with α* = 0 (***K***_0_*) and* 0.5 (***K***_0.5_*) are compared.*

| Simulation Design | Method | Kernel Type |  |  |  |  |
| --- | --- | --- | --- | --- | --- | --- |
| | | $\mathbf{K}_W$ | $\mathbf{K}_U$ | $\mathbf{K}_{VAW}$ | $\mathbf{K}_0$ | $\mathbf{K}_{0.5}$ |
| Cross-sectional | eRLRT | 0.046 | 0.043 | 0.045 | 0.048 | 0.047 |
|  | eLRT | 0.046 | 0.043 | 0.051 | 0.052 | 0.046 |
|  | eScore | 0.039 | 0.031 | 0.047 | 0.045 | 0.042 |
| Longitudinal | eRLRT | 0.041 | 0.053 | 0.045 | 0.041 | 0.042 |
|  | eScore | 0.034 | 0.048 | 0.048 | 0.050 | 0.045 |

**Figure 1.**
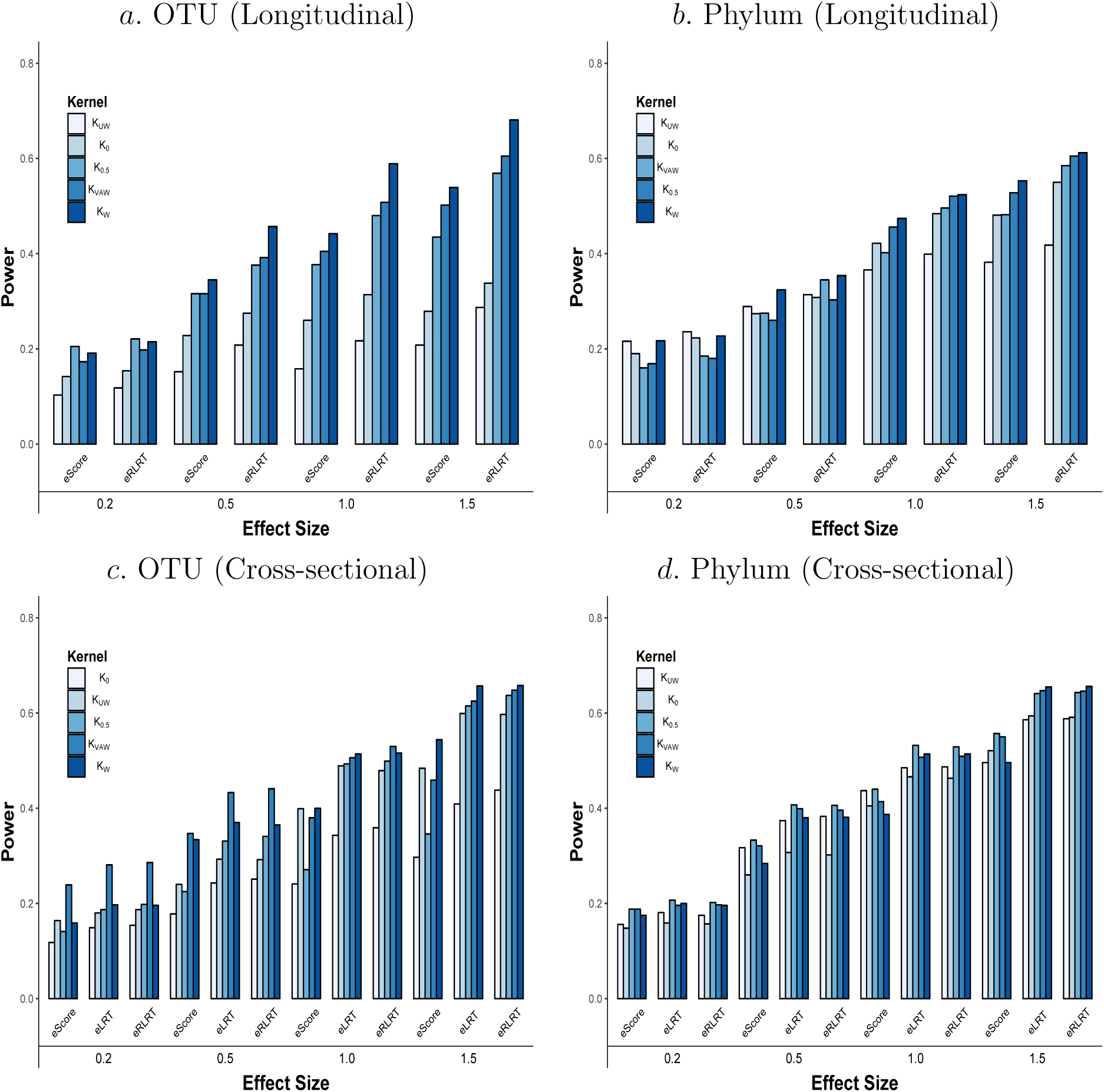
Scenario 1: Power of eRLRT, eLRT and eScore using different distance measures. Left figure shows results where the OTU counts are used to calculate distances, right figures shows that OTU counts are summarized at phylum level to construct the distances. *K*_0_, *K*_0.5_, *K*_*W*_, *K*_*U*_ and *K*_*V* *AW*_ represent generalized UniFrac distance with *α* = 0, 0.5, weighted UniFrac distance, unweighted UniFrac distance and variance adjusted weighted UniFrac distance, respectively.

**Scenario 2. Localizing fine microbiome cluster effects.** Table 3 shows the type I error rates for testing microbiome effect at the phylum level, with and without adjusting for the effect contributed by cluster, *other*. Most of the type I error rates are inflated when not adjusting for cluster *other* effects. In cross-sectional design, the type I error rates of *Bacteroidetes* and *Proteobacteria* stay correct due to its weak correlation with cluster *other* (Pearson correlation = 0.04, 0.11 with *p*-value = 0.70, 0.24, respectively). After adjustment, type I error rates stay correct even when confounding effects are large (Table 3).

**Table 3.**
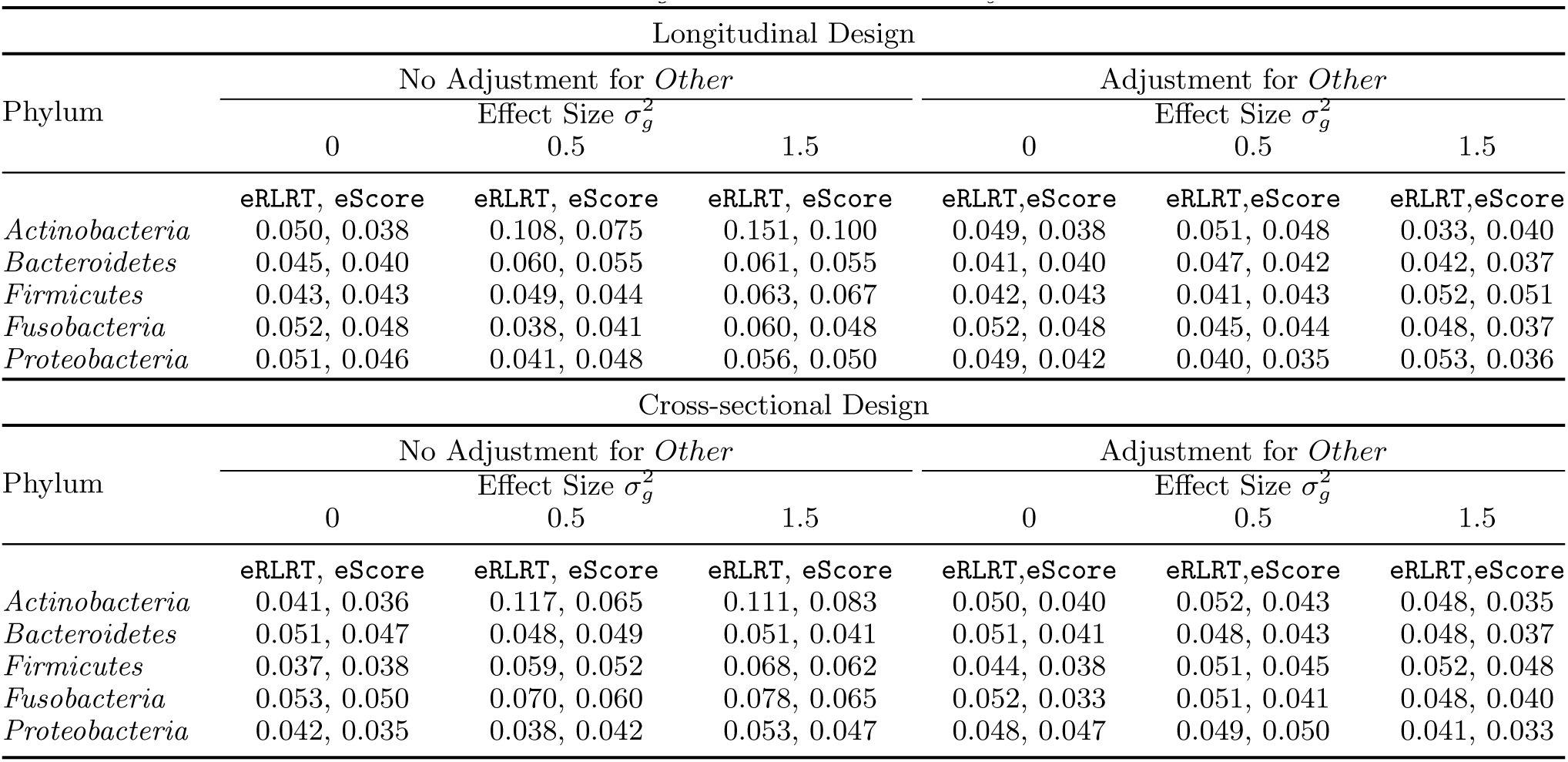
*Scenario 2: Type I error rate of localizing fine microbiome cluster effects. Only cluster “Other” contains effects*, 0, 0.5 *and* 1.5. *Type I error rates are evaluated with or without adjustment for effect from cluster Other. The weighted UniFrac kernel is used. Top panel shows results from simulation using longitudinal data while bottom panel shows results using cross-sectional data only.*

| Longitudinal Design |  |  |  |  |  |  |
| --- | --- | --- | --- | --- | --- | --- |
| Phylum | No Adjustment for *Other* |  |  | Adjustment for *Other* |  |  |
| | Effect Size $\sigma_g^2$ | | | Effect Size $\sigma_g^2$ | | |
|  | 0 | 0.5 | 1.5 | 0 | 0.5 | 1.5 |
|  | eRLRT, eScore | eRLRT, eScore | eRLRT, eScore | eRLRT,eScore | eRLRT,eScore | eRLRT,eScore |
| *Actinobacteria* | 0.050, 0.038 | 0.108, 0.075 | 0.151, 0.100 | 0.049, 0.038 | 0.051, 0.048 | 0.033, 0.040 |
| *Bacteroidetes* | 0.045, 0.040 | 0.060, 0.055 | 0.061, 0.055 | 0.041, 0.040 | 0.047, 0.042 | 0.042, 0.037 |
| *Firmicutes* | 0.043, 0.043 | 0.049, 0.044 | 0.063, 0.067 | 0.042, 0.043 | 0.041, 0.043 | 0.052, 0.051 |
| *Fusobacteria* | 0.052, 0.048 | 0.038, 0.041 | 0.060, 0.048 | 0.052, 0.048 | 0.045, 0.044 | 0.048, 0.037 |
| *Proteobacteria* | 0.051, 0.046 | 0.041, 0.048 | 0.056, 0.050 | 0.049, 0.042 | 0.040, 0.035 | 0.053, 0.036 |
| Cross-sectional Design |  |  |  |  |  |  |
| Phylum | No Adjustment for *Other* |  |  | Adjustment for *Other* |  |  |
| | Effect Size $\sigma_g^2$ | | | Effect Size $\sigma_g^2$ | | |
|  | 0 | 0.5 | 1.5 | 0 | 0.5 | 1.5 |
|  | eRLRT, eScore | eRLRT, eScore | eRLRT, eScore | eRLRT,eScore | eRLRT,eScore | eRLRT,eScore |
| *Actinobacteria* | 0.041, 0.036 | 0.117, 0.065 | 0.111, 0.083 | 0.050, 0.040 | 0.052, 0.043 | 0.048, 0.035 |
| *Bacteroidetes* | 0.051, 0.047 | 0.048, 0.049 | 0.051, 0.041 | 0.051, 0.041 | 0.048, 0.043 | 0.048, 0.037 |
| *Firmicutes* | 0.037, 0.038 | 0.059, 0.052 | 0.068, 0.062 | 0.044, 0.038 | 0.051, 0.045 | 0.052, 0.048 |
| *Fusobacteria* | 0.053, 0.050 | 0.070, 0.060 | 0.078, 0.065 | 0.052, 0.033 | 0.051, 0.041 | 0.048, 0.040 |
| *Proteobacteria* | 0.042, 0.035 | 0.038, 0.042 | 0.053, 0.047 | 0.048, 0.047 | 0.049, 0.050 | 0.041, 0.033 |

In practice symbiosis of bacteria causes correlation between them (Xu et al., 2007; Dickson et al., 2013; Zeng et al., 2016). Medication targeting of specific pathogens can minimize damage to essential symbiotic microbial species, and preserve community structure and function in the healthy (and developing) microbiome (Hicks et al., 2013; Blaser, 2016). Simulation scenario 2 demonstrates that our method is equipped to localizing fine microbiome cluster effects.

**Scenario 3: Comparing to existing methods** `MiRKAT` **and** `LinScore`. Table 4 presents the type I error rate and power for `eRLRT`, `eLRT`, `eScore`, `MiRKAT` and `LinScore` tests in detecting overall microbiome effects. The power is presented for both cross-sectional and longitudinal studies with sample size from 20 to 100. `eRLRT` and `eLRT` outperform `LinScore` and `MiRKAT` in baseline simulation studies. For simulation with repeated measurements, `eRLRT` outperforms `LinScore` under small sample sizes. For sample size *n* = 100, `eRLRT` has similar or slightly higher power comparing to `LinScore` when association strength is weak. Microbiome studies usually have limited sample size due to the high cost. Higher power of the exact tests at small sample sizes will be particularly valuable for biologists and physicians to identify the associated microbiome clusters.

**Table 4.** *Scenario 3: Comparing to existing methods. Type I error rate and power from `eLRT`, `eRLRT`, `eScore`, `LinScore`, and `MiRKAT` at baseline when* #*Repeat* = 1. *When* #*Repeat* = 2, *only `LinScore` is compared with `eRLRT` and `eScore`. Sample sizes (n) range from* 20 *to* 100 *and effect sizes* 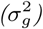 *range from* 0 *to* 1.5.

| $n$ | #Repeat | Method | Effect Size ($\sigma_g^2$) | | | | | | |
| --- | --- | --- | --- | --- | --- | --- | --- | --- | --- |
|  |  |  | 0 | 0.10 | 0.2 | 0.5 | 0.8 | 1.0 | 1.5 |
| 20 | 1 | *eScore* | 0.045 | 0.059 | 0.050 | 0.074 | 0.078 | 0.079 | 0.104 |
|  |  | *eLRT* | 0.051 | 0.089 | 0.095 | 0.111 | 0.118 | 0.141 | 0.152 |
|  |  | *eRLRT* | 0.050 | 0.097 | 0.088 | 0.108 | 0.122 | 0.142 | 0.160 |
|  |  | *MiRKAT* | 0.048 | 0.056 | 0.046 | 0.071 | 0.069 | 0.077 | 0.104 |
|  |  | *LinScore* | 0.050 | 0.060 | 0.046 | 0.075 | 0.072 | 0.078 | 0.106 |
|  | 2 | *eScore* | 0.050 | 0.055 | 0.040 | 0.057 | 0.068 | 0.077 | 0.088 |
|  |  | *eRLRT* | 0.051 | 0.055 | 0.074 | 0.081 | 0.092 | 0.085 | 0.118 |
|  |  | *LinScore* | 0.049 | 0.057 | 0.063 | 0.055 | 0.072 | 0.078 | 0.090 |
| 30 | 1 | *eScore* | 0.043 | 0.059 | 0.050 | 0.074 | 0.078 | 0.079 | 0.104 |
|  |  | *eLRT* | 0.046 | 0.089 | 0.095 | 0.111 | 0.118 | 0.141 | 0.152 |
|  |  | *eRLRT* | 0.052 | 0.097 | 0.088 | 0.108 | 0.122 | 0.142 | 0.160 |
|  |  | *MiRKAT* | 0.055 | 0.056 | 0.046 | 0.071 | 0.069 | 0.077 | 0.104 |
|  |  | *LinScore* | 0.054 | 0.060 | 0.046 | 0.075 | 0.072 | 0.078 | 0.106 |
|  | 2 | *eScore* | 0.045 | 0.058 | 0.067 | 0.093 | 0.114 | 0.127 | 0.151 |
|  |  | *eRLRT* | 0.052 | 0.063 | 0.081 | 0.105 | 0.127 | 0.145 | 0.178 |
|  |  | *LinScore* | 0.046 | 0.054 | 0.061 | 0.076 | 0.088 | 0.132 | 0.134 |
| 50 | 1 | *eScore* | 0.036 | 0.070 | 0.071 | 0.118 | 0.151 | 0.164 | 0.240 |
|  |  | *eLRT* | 0.048 | 0.084 | 0.094 | 0.135 | 0.188 | 0.214 | 0.306 |
|  |  | *eRLRT* | 0.049 | 0.086 | 0.088 | 0.127 | 0.192 | 0.201 | 0.307 |
|  |  | *MiRKAT* | 0.047 | 0.065 | 0.069 | 0.114 | 0.156 | 0.183 | 0.257 |
|  |  | *LinScore* | 0.045 | 0.070 | 0.077 | 0.124 | 0.176 | 0.189 | 0.267 |
|  | 2 | *eScore* | 0.047 | 0.069 | 0.084 | 0.110 | 0.148 | 0.177 | 0.257 |
|  |  | *eRLRT* | 0.041 | 0.074 | 0.097 | 0.134 | 0.188 | 0.217 | 0.315 |
|  |  | *LinScore* | 0.051 | 0.063 | 0.096 | 0.156 | 0.205 | 0.261 | 0.333 |
| 100 | 1 | *eScore* | 0.050 | 0.096 | 0.165 | 0.304 | 0.383 | 0.390 | 0.532 |
|  |  | *eLRT* | 0.052 | 0.114 | 0.191 | 0.377 | 0.472 | 0.516 | 0.664 |
|  |  | *eRLRT* | 0.049 | 0.105 | 0.195 | 0.375 | 0.460 | 0.510 | 0.661 |
|  |  | *MiRKAT* | 0.051 | 0.093 | 0.181 | 0.329 | 0.427 | 0.483 | 0.622 |
|  |  | *LinScore* | 0.048 | 0.106 | 0.194 | 0.347 | 0.439 | 0.507 | 0.630 |
|  | 2 | *eScore* | 0.037 | 0.140 | 0.205 | 0.277 | 0.378 | 0.411 | 0.525 |
|  |  | *eRLRT* | 0.041 | 0.161 | 0.244 | 0.327 | 0.447 | 0.498 | 0.626 |
|  |  | *LinScore* | 0.046 | 0.121 | 0.214 | 0.347 | 0.451 | 0.545 | 0.652 |

### Analysis of Longitudinal Pulmonary Microbiome Data

It is known that HIV infection is associated with alterations in the respiratory microbiome (Twigg III et al., 2016). However due to the limited investigation, the clinical implications of lung microbial dysbiosis are currently unknown. As the initial step to reveal the mechanistic of respiratory microbiome to pulmonary complications in HIV-infected individuals, we investigate the relationship between pulmonary function and the respiratory microbiota profiles in the bronchoalveolar lavage (BAL) fluid of 30 HIV-infected patients at the advanced stage (baseline mean CD4 count, 262 cells/*mm*^3^). Their acellular BAL fluid was sampled at baseline, 4 weeks, 1 year, and 3 years. 16S rRNA gene sequencing technology was used to study pulmonary microbiota. Details of microbiome composition has been discussed in Section 3. Pulmonary function is measured by spirometry and diffusion capacity tests. Spirometry tests measure how much and how quickly air can move out of lung. Typical spirometry tests include forced vital capacity (FVC), forced expiratory volume in 1 second (FEV1), and average forced expiratory flow (FEF). Diffusion capacity of the lungs for carbon monoxide (DLCO) measures how much oxygen travels from the alveoli of the lungs to the blood stream. DLCO corrected for hemoglobin (DsbHb) and diffusion capacity corrected for alveolar volume and hemoglobin (DVAsbHb) are evaluated. Descriptive statistics of these measures are summarized in Web Appendix Table 1.

Exact tests and `LinScore` are used to study the association. Associations with p-values less than 0.05 are reported to be significant. Covariates include gender, race, smoking status, CD4 counts, and HIV virus load (Table 5). Missing covariate is imputed by its mean. For overall microbiome association test, no tests find significant associations. However at the phylum level, *Bacteroidetes* shows significant association with spirometry while *Firmicutes* shows significant association with diffusing capacity measures. Similar results have previously been reported by Tunney et al. (2013); Molyneaux et al. (2012). We then focus on analyzing genera from both phyla *Bacteroidetes* and *Firmicutes* given their important status in normal lungs (Cui et al., 2014). Only by `eRLRT` and `eScore`, genus *Prevotella*, *Porphyromonas*, and *Parvimonas* show significant effects on FEF and FEV1 (Table 5). Genus *Veillonella* shows significant association with FEF. It appears that both *Parvimonas* and *Veillonella* in phylum *Firmicutes* are significantly associated with FEF and both genus *Prevotella* and *Porphyromonas* in phylum *Bacteroidetes* are significantly associated with FEF and FEV1. We therefore perform the test in a joint model to localize fine cluster effect. Interestingly, by `eRLRT` the significant association between genus *Parvimonas* and FEF still remains after adjusting for the effects from genus *Veillonella*. But the opposite is not true. This supports the previous studies that *Parvimonas* abundance changed in subjects with pulmonary disease (e.g., asthma or COPD) comparing to the control group (Pragman et al., 2012; Kim et al., 2017). However, either *Prevotella* or *Porphyromonas* lost its significance when adjusting for the other. This likely suggests that *Prevotella* and *Porphyromonas* are correlated and both tag effects to lung function. In comparison, `LinScore` only detects the significant microbiome effect of *Bacteroidetes* with FEF. Our results further support the conclusions from previous studies and shed lights for future clinical causality research (Twigg III et al., 2016; Weiden et al., 2017; Segal et al., 2017). None of the tests (exact tests, `LinScore`, and `MiRKAT`) identify significant associations using only baseline data (results not shown).

**Table 5.**
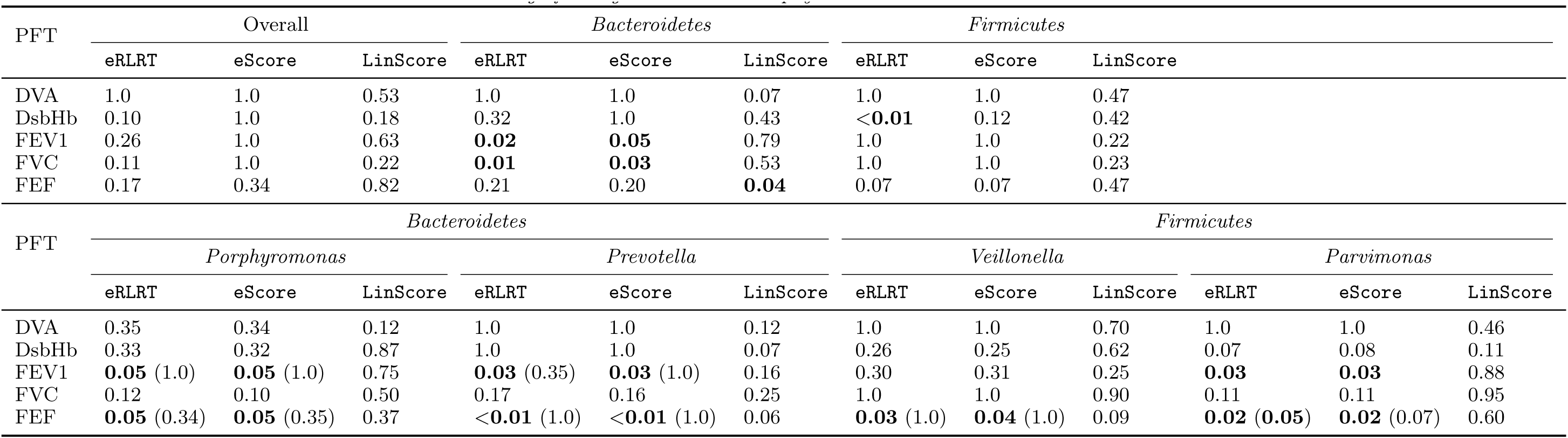
*Application to the longitudinal pulmonary microbiome studies. `eRLRT`, `eScore`, and `LinScore` are used to detect association. Genus Porphyromonas and Prevotella belong to phylum Bacteroidetes while genus Veillonella and Parvimonas belong to phylum Firmicutes. Upper panel shows the testing results at phylum level, while lower panel shows the results at genus level. P-values less than* 0.05 *are highlighted in bold font. P-values in parenthesis show the results from a joint model where significant genus in the same phylum are included.*

## 5. Discussion

In this report, motivated by a longitudinal pulmonary microbiome study, we develop and implement three computationally efficient exact variance component tests (`eScore`, `eLRT`, and `eRLRT`). Our method extend previous exact variance component tests to the case when the null hypothesis contains more than one variance component (Zhou et al., 2016). They can be applied to longitudinal studies testing the overall microbiome effects, as well as crosssectional studies identifying microbiome associations at fine-grained level. The latter has been emerging as the focus of many current microbiome studies (Nayfach et al., 2016; Lloyd-Price et al., 2017; Truong et al., 2017). Unlike Qu et al. (2013) and Zhao et al. (2015)’s score test that uses moment-matching to approximate null distribution, our tests are exact in finite samples, therefore beneficial to the studies with limited sample size. Compared to score test, our `eLRT` and `eRLRT` tests can further boost testing power when microbiome effects are weak. Simulation studies verify that our exact tests have correct size and many innovative utilizations. In the application to the real longitudinal pulmonary microbiome study, only our exact tests detect multiple interesting genera associated with lung function. We then further demonstrate the ability of our exact tests to differentiate associated genus by two real data examples. Although the derivation of `eLRT` and `eRLRT` require normality assumption, a sensitivity simulation shows that even with a misspecified phenotypic distribution, like *t*distribution, our tests still preserve correct Type I error rate (Web Appendix E, Table 2). The software package is implemented in the open source, high-performance technical computing language Julia and is freely available. We also offer many types of distance calculation to further ease the computation and advance microbiome studies.

There are a few directions for future work. First, there are linear mixed effects models not of form (3), for example, those include both random intercepts and random slopes (Drikvandi et al., 2013). Our methods extend to these cases naturally and we defer them to future research. Second direction is to incorporate multiple types of kernels into exact tests. Last we consider extension to the generalized linear mixed effects models, although it can be challenging especially for LRT and RLRT. Score-based tests may be possible through penalized quasi-likelihood (PQL) (Lin, 1997; Chen et al., 2016).

## 6. Software

A Julia package is freely available at https://github.com/JingZhai63/VCmicrobiome.

## 7. Supplementary Materials

Web Appendix, Tables, and Figures, referenced in the paper are available at the Biometrics website on Wiley Online Library.

## Acknowledgments

JJZ is supported by NIH grant K01DK106116 and Arizona Biomedical Research Commission (ABRC) grant. HZ is partially supported by NIH grants HG006139, GM105785, GM53275 and NSF grant DMS-1645093.

## References

Altschul, S. F., Gish, W., Miller, W., Myers, E. W., and Lipman, D. J. (1990). Basic local alignment search tool. Journal of Molecular Biology 215, 403 – 410.

Blaser, M. J. (2016). Antibiotic use and its consequences for the normal microbiome. Science 352, 544–545.

BLAST (1990). BLAST: Basic Local Alignment Search Tool. https://blast.ncbi.nlm.nih.gov/Blast.cgi. (Accessed 2018-01-11).

Bray, J. R. and Curtis, J. T. (1957). An ordination of the upland forest communities of southern Wisconsin. Ecological Monographs 27, 325–349.

Chang, Q., Luan, Y., and Sun, F. (2011). Variance adjusted weighted UniFrac: a powerful beta diversity measure for comparing communities based on phylogeny. BMC Bioinformatics 12, 118.

Chen, E. Z. and Li, H. (2016). A two-part mixed-effects model for analyzing longitudinal microbiome compositional data. Bioinformatics 32, 2611–2617.

Chen, H., Wang, C., Conomos, M. P., Stilp, A. M., Li, Z., Sofer, T., Szpiro, A. A., Chen, W., Brehm, J. M., Celedón, J. C., et al. (2016). Control for population structure and relatedness for binary traits in genetic association studies via logistic mixed models. The American Journal of Human Genetics 98, 653–666.

Chen, J., Bittinger, K., Charlson, E. S., Hoffmann, C., Lewis, J., Wu, G. D., Collman, R. G., Bushman, F. D., and Li, H. (2012). Associating microbiome composition with environmental covariates using generalized UniFrac distances. Bioinformatics 28, 2106–2113.

Chen, J., Chen, W., Zhao, N., Wu, M. C., and Schaid, D. J. (2016). Small sample kernel association tests for human genetic and microbiome association studies. Genetic Epidemiology 40, 5–19.

Chen, Z. and Dunson, D. B. (2003). Random effects selection in linear mixed models. Biometrics 59, 762–769.

Christensen, R. (1996). Exact tests for variance components. Biometrics 52, 309–314.

Crainiceanu, C. M. and Ruppert, D. (2004). Likelihood ratio tests in linear mixed models with one variance component. Journal of the Royal Statistical Society: Series B (Statistical Methodology) 66, 165–185.

Cui, L., Morris, A., Huang, L., Beck, J. M., Twigg III, H. L., Von Mutius, E., and Ghedin, E. (2014). The microbiome and the lung. Annals of the American Thoracic Society 11, S227–S232.

Dickson, R. P., Erb-Downward, J. R., and Huffnagle, G. B. (2013). The role of the bacterial microbiome in lung disease. Expert Review of Respiratory Medicine 7, 245–257.

Drik-vandi, R., Verbeke, G., Khodadadi, A., and Partovi Nia, V. (2013). Testing multiple variance components in linear mixed-effects models. Biostatistics 14, 144–59.

Eckburg, P. B., Bik, E. M., Bernstein, C. N., Purdom, E., Dethlefsen, L., Sargent, M., Gill, S. R., Nelson, K. E., and Relman, D. A. (2005). Diversity of the human intestinal microbial flora. Science 308, 1635–1638.

Erb-Downward, J. R., Thompson, D. L., Han, M. K., Freeman, C. M., McCloskey, L., Schmidt, L. A., Young, V. B., Toews, G. B., Curtis, J. L., Sundaram, B., et al. (2011). Analysis of the lung microbiome in the “healthy” smoker and in COPD. PloS One 6, e16384.

Federhen, S. (2012). The NCBI Taxonomy database. Nucleic Acids Research 40, D136–D143.

Gilbert, J. A., Quinn, R. A., Debelius, J., Xu, Z. Z., Morton, J., Garg, N., Jansson, J. K., Dorrestein, P. C., and Knight, R. (2016). Microbiome-wide association studies link dynamic microbial consortia to disease. Nature 535, 94–103.

Greven, S., Crainiceanu, C. M., Küchenhoff, H., and Peters, A. (2008). Restricted likelihood ratio testing for zero variance components in linear mixed models. Journal of Computational and Graphical Statistics 17, 870–891.

Grubb, J. R., Moorman, A. C., Baker, R. K., Masur, H., et al. (2006). The changing spectrum of pulmonary disease in patients with HIV infection on antiretroviral therapy. Aids 20, 1095–1107.

Haas, B. J., Gevers, D., Earl, A. M., Feldgarden, M., Ward, D. V., Giannoukos, G., Ciulla, D., Tabbaa, D., Highlander, S. K., Sodergren, E., et al. (2011). Chimeric 16s rRNA sequence formation and detection in Sanger and 454-pyrosequenced PCR amplicons. Genome Research 21, 494–504.

Hicks, L. A., Taylor Jr, T. H., and Hunkler, R. J. (2013). US outpatient antibiotic prescribing, 2010. New England Journal of Medicine 368, 1461–1462.

Hodkinson, B. P. and Grice, E. A. (2015). Next-generation sequencing: a review of technologies and tools for wound microbiome research. Advances in Wound Care 4, 50–58.

Jangi, S., Gandhi, R., Cox, L. M., Li, N., Von Glehn, F., Yan, R., Patel, B., Mazzola, M. A., Liu, S., Glanz, B. L., et al. (2016). Alterations of the human gut microbiome in multiple sclerosis. Nature Communications 7, 12015.

Kim, B.-S., Lee, E., Lee, M.-J., Kang, M.-J., Yoon, J., Cho, H.-J., Park, J., Won, S., Lee, Y., and Hong, S. J. (2017). Different functional genes of upper airway microbiome S. associated with natural course of childhood asthma. Allergy.

Kinney, S. K. and Dunson, D. B. (2007). Fixed and random effects selection in linear and logistic models. Biometrics 63, 690–698.

Kuleshov, V., Jiang, C., Zhou, W., Jahanbani, F., Batzoglou, S., and Snyder, M. (2016). Synthetic long-read sequencing reveals intraspecies diversity in the human microbiome. Nature Biotechnology 34, 64–69.

Letunic, I. and Bork, P. (2007). Interactive Tree of Life (iTOL): an online tool for phylogenetic tree display and annotation. Bioinformatics 23, 127–128.

Letunic, I. and Bork, P. (2011). Interactive Tree of Life v2: online annotation and display of phylogenetic trees made easy. Nucleic Acids Research 39, W475–W478.

Lin, X. (1997). Variance component testing in generalised linear models with random effects. Biometrika 84, 309–326.

Lloyd-Price, J., Mahurkar, A., Rahnavard, G., Crabtree, J., Orvis, J., Hall, A. B., Brady, A., Creasy, H. H., McCracken, C., Giglio, M. G., et al. (2017). Strains, functions and dynamics in the expanded human microbiome project. Nature 550, 61.

Lozupone, C., Cota-Gomez, A., Palmer, B. E., Linderman, D. J., Charlson, E. S., Sodergren, E., Mitreva, M., Abubucker, S., Martin, J., Yao, G., et al. (2013). Widespread colonization of the lung by Tropheryma whipplei in HIV infection. American Journal of Respiratory and Critical Care Medicine 187, 1110–1117.

Lozupone, C. and Knight, R. (2005). UniFrac: a new phylogenetic method for comparing microbial communities. Applied and Environmental Microbiology 71, 8228–8235.

Lozupone, C. A., Hamady, M., Kelley, S. T., and Knight, R. (2007). Quantitative and qualitative *β* diversity measures lead to different insights into factors that structure microbial communities. Applied and Environmental Microbiology 73, 1576–1585.

Maidak, B. L., Olsen, G. J., Larsen, N., Overbeek, R., McCaughey, M. J., and Woese, C. R. T. (1996). The Ribosomal Database Project (RDP). Nucleic Acids Research 24, 82–85.

Molyneaux, P. L., Russell, A. M., Cox, M. J., Moffatt, M. F., Cookson, W. O., and Maher, M. (2012). The respiratory microbiome in idiopathic pulmonary fibrosis. In C103. pathogenesis, biomarkers, and risk factors for interstitial lung disease: from bench to bedside, pages A5174–A5174. American Thoracic Society.

S. Morris, A., Beck, J. M., Schloss, P. D., Campbell, T. B., Crothers, K., Curtis, J. L., Flores, C., Fontenot, A. P., Ghedin, E., Huang, L., et al. (2013). Comparison of the respiratory microbiome in healthy nonsmokers and smokers. American Journal of Respiratory and Critical Care Medicine 187, 1067–1075.

Mothur (2009). Mothur: A software for describing and comparing microbial communities. https://www.mothur.org. (Accessed 2018-01-11).

Nayfach, S., Rodriguez-Mueller, B., Garud, N., and Pollard, K. S. (2016). An integrated metagenomics pipeline for strain profiling reveals novel patterns of bacterial transmission and biogeography. Genome research 26, 1612–1625.

Ofversten, J. (1993). Exact tests for variance components in unbalanced mixed linear models. Biometrics 49, 45–57.

PhylogeneticDistance.jl (2017). PhylogeneticDistance.jl: A julia package for calculating phylogenetic distance. https://github.com/JingZhai63/PhylogeneticDistance.jl. (Accessed 2018-01-11).

phyloT (2006). phyloT: A phylogenetic tree generator. http://phylot.biobyte.de/. (Accessed 2018-01-11).

Pragman, A. A., Kim, H. B., Reilly, C. S., Wendt, C., and Isaacson, R. E. (2012). The lung microbiome in moderate and severe chronic obstructive pulmonary disease. PloS one 7, e47305.

Qu, L., Guennel, T., and Marshall, S. L. (2013). Linear score tests for variance components in linear mixed models and applications to genetic association studies. Biometrics 69, 883–892.

Rogers, G. B., Carroll, M., Hoffman, L., Walker, A., Fine, D., and Bruce, K. (2010). Comparing the microbiota of the cystic fibrosis lung and human gut. Gut Microbes 1, 85–93.

Saville, B. R. and Herring, A. H. (2009). Testing random effects in the linear mixed model using approximate bayes factors. Biometrics 65, 369–76.

Schloss, P. D., Westcott, S. L., Ryabin, T., Hall, J. R., Hartmann, M., Hollister, E. B., Lesniewski, R. A., Oakley, B. B., Parks, D. H., Robinson, C. J., et al. (2009). Introducing mothur: open-source, platform-independent, community-supported software for describing and comparing microbial communities. Applied and Environmental Microbiology 75, 7537–7541.

Segal, L. N., Clemente, J. C., Li, Y., Ruan, C., Cao, J., Danckers, M., Morris, A., Tapyrik, S., Wu, B. G., Diaz, P., et al. (2017). Anaerobic bacterial fermentation products increase tuberculosis risk in antiretroviral-drug-treated HIV patients. Cell Host & Microbe 21, 530–537.

Self, S. G. and Liang, K.-Y. (1987). Asymptotic properties of maximum likelihood estimators and likelihood ratio tests under nonstandard conditions. Journal of the American Statistical Association 82, 605–610.

Silvapulle, M. J. and Sen, P. K. (2011). Constrained statistical inference: order, inequality, and shape constraints, volume 912.

Stram, D. O. and Lee, J. W. (1994). Variance components testing in the longitudinal mixed effects model. Biometrics pages 1171–1177.

Taxonomy (1991). NCBI taxonomy database. https://www.ncbi.nlm.nih.gov/taxonomy. (Accessed 2018-01-11).

Truong, D. T., Tett, A., Pasolli, E., Huttenhower, C., and Segata, N. (2017). Microbial strain-level population structure and genetic diversity from metagenomes. Genome research 27, 626–638.

Tunney, M. M., Einarsson, G. G., Wei, L., Drain, M., Klem, E. R., Cardwell, C., Ennis, M., Boucher, R. C., Wolfgang, M. C., and Elborn, J. S. (2013). Lung microbiota and bacterial abundance in patients with bronchiectasis when clinically stable and during exacerbation. American Journal of Respiratory and Critical Care Medicine 187, 1118–1126.

T. Twigg, H. L., Morris, A., Ghedin, E., Curtis, J. L., Huffnagle, G. B., Crothers, K., Campbell, B., Flores, S. C., Fontenot, A. P., Beck, J. M., et al. (2013). Use of bronchoalveolar lavage to assess the respiratory microbiome: signal in the noise. The Lancet Respiratory Medicine 1, 354–356.

Twigg III, H. L., Knox, K. S., Zhou, J., Crothers, K. A., Nelson, D. E., Toh, E., Day, R. B., Lin, H., Gao, X., Dong, Q., et al. (2016). Effect of advanced HIV infection on the respiratory microbiome. American Journal of Respiratory and Critical Care Medicine 194, 226–235.

Wang, J. and Jia, H. (2016). Metagenome-wide association studies: fine-mining the microbiome. Nature Reviews Microbiology 14, 508–522.

Weiden, M. D., Segal, L. N., Clemente, J., Li, Y., Danckers-Degregory, M., Morris, A. M., Tapyrik, S., Diaz, P., Dawson, R., Van Zyl-Smit, R., et al. (2017). Lung microbiome dysbiosis is a risk factor for pulmonary diffusion abnormalities in antiretroviral treated HIV-infection. In A13. Role of dysbiosis in lung disease, pages A1002–A1002. American Thoracic Society.

Xu, J., Mahowald, M. A., Ley, R. E., Lozupone, C. A., Hamady, M., Martens, E. C., Henrissat, B., Coutinho, P. M., Minx, P., Latreille, P., et al. (2007). Evolution of symbiotic bacteria in the distal human intestine. PLoS Biology 5, 1–13.

Zemanick, E. T., Sagel, S. D., and Harris, J. K. (2011). The airway microbiome in cystic fibrosis and implications for treatment. Current Opinion in Pediatrics 23, 319–324.

Zeng, M. Y., Cisalpino, D., Varadarajan, S., Hellman, J., Warren, H. S., Cascalho, M., Inohara, N., and Núñez, G. (2016). Gut microbiota-induced immunoglobulin G controls systemic infection by symbiotic bacteria and pathogens. Immunity 44, 647–658.

Zhan, X., Tong, X., Zhao, N., Maity, A., Wu, M. C., and Chen, J. (2017). A small-sample multivariate kernel machine test for microbiome association studies. Genetic Epidemiology 41, 210–220.

Zhao, N., Chen, J., Carroll, I. M., Ringel-Kulka, T., Epstein, M. P., Zhou, H., Zhou, J. J., Ringel, Y., Li, H., and Wu, M. C. (2015). Testing in microbiome-profiling studies with MiRKAT, the microbiome regression-based kernel association test. American Journal of Human Genetics 96, 797–807.

Zhou, J. J., Hu, T., Qiao, D., Cho, M. H., and Zhou, H. (2016). Boosting gene mapping power and efficiency with efficient exact variance component tests of single nucleotide polymorphism sets. Genetics 204, 921–931.

